## Supplementary Materials for "Resolving the conformational ensemble of a membrane protein by integrating small-angle scattering with AlphaFold"

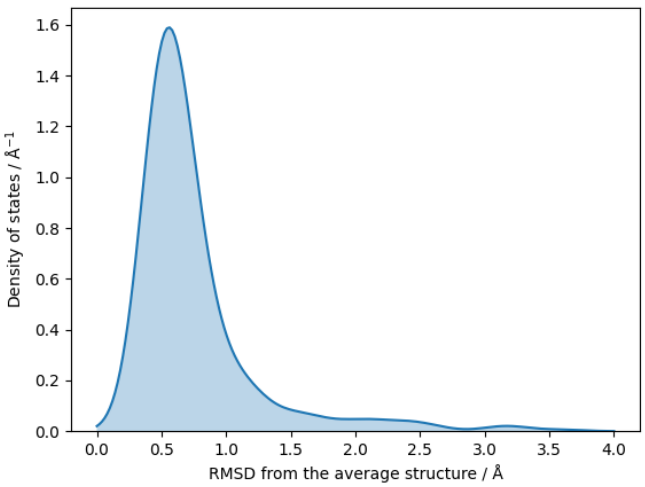


**S1 Fig. Variability in the AF-sampled ensemble.** Probability density of the root mean square deviation of the C$\alpha$ atoms of the AF-generated conformations with pLDDT $\geq 75$, compared to the average structure. Alignment is done on the C$\alpha$ atoms. Two outliers (with RMSDs of 6.9 Å and 9.0 Å respectively) are not shown.

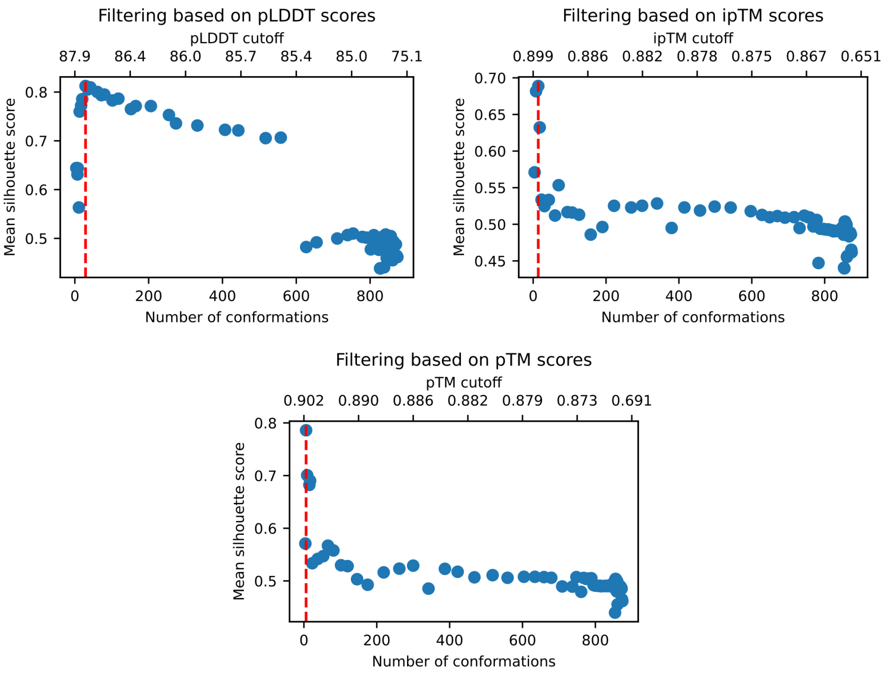


**S3 Fig. Clustering quality for different AF-quality metrics.** Average silhouette score of the agglomerative clustering of the SANS intensity profiles of all conformations with pLDDT (top left), ipTM (top right), or pTM (bottom) scores above different cutoffs for the initial run of the pipeline, as a function of the number of such conformations. The dashed red line indicates the cutoff for the maximal silhouette score.

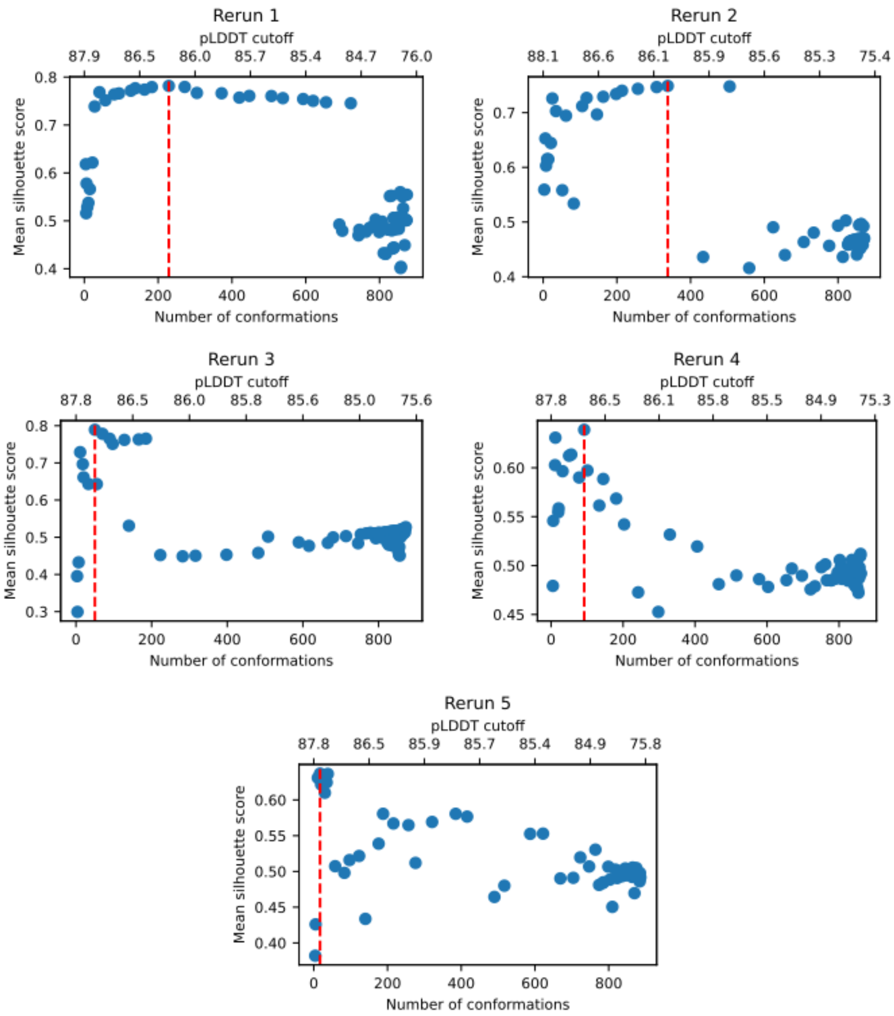


**S4 Fig. Clustering quality across the reruns of the pipeline.** Average silhouette score of the agglomerative clustering of the SANS intensity profiles of all conformations with pLDDT scores above different cutoffs for the five reruns, as a function of the number of such conformations. The dashed red line indicates the cutoff for the maximal silhouette score.

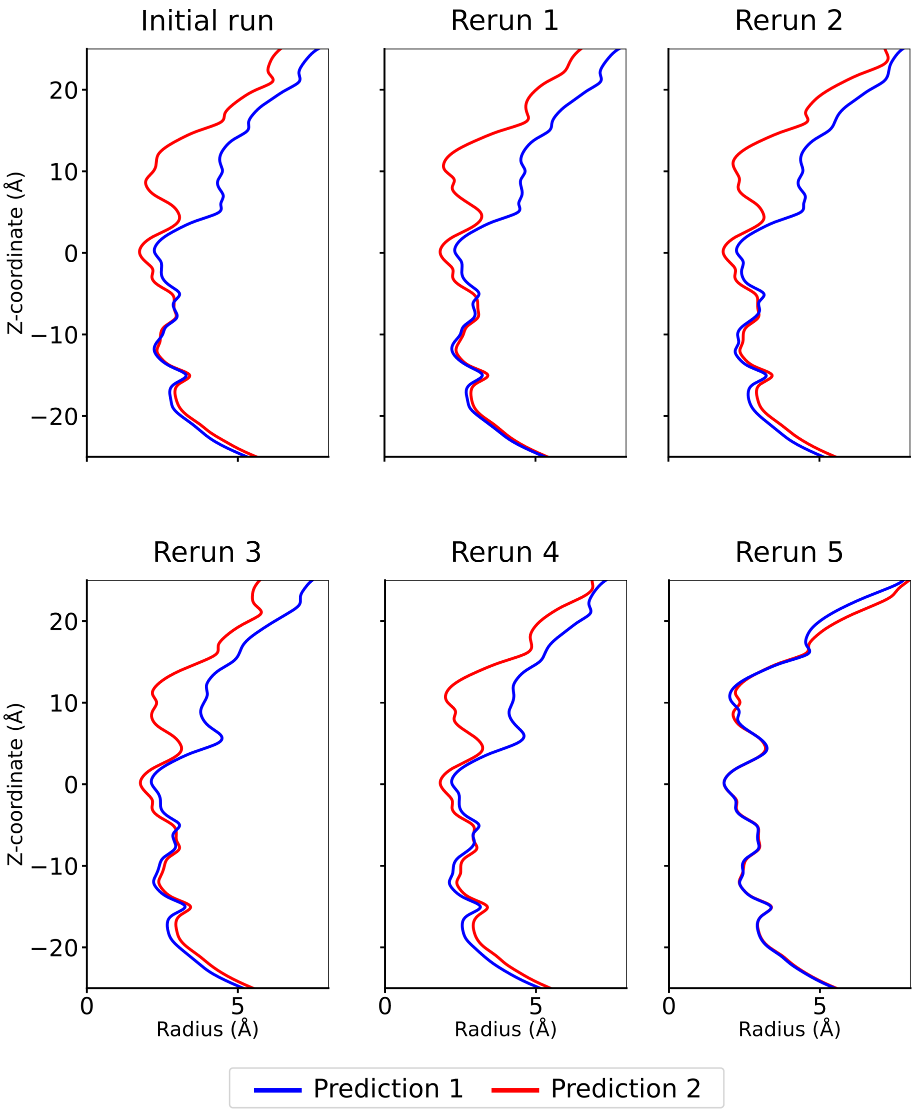


**S5 Fig. Pore profiles of the selected conformations.**

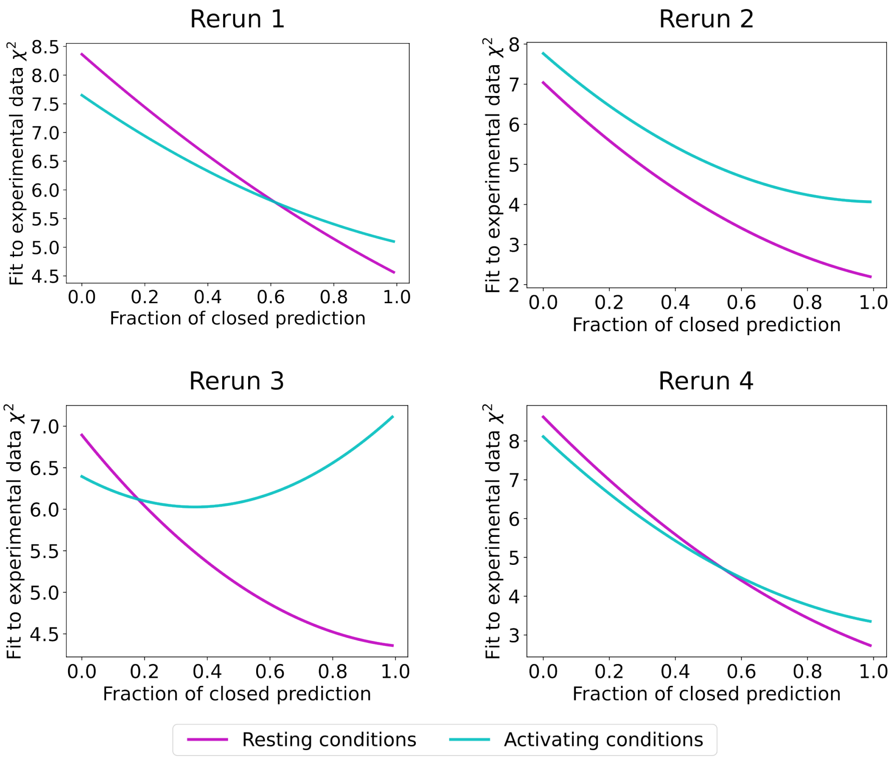


**S6 Fig. Fits of selected conformations to experimental SANS data.** Fits to the experimental SANS data for a linear combination of the closed and open prediction as a function of their relative weights, for the four reruns in which distinct functional states were predicted.

**S7 Table Cα RMSD of the predicted structures compared to experimental crystal structures.**

| Data set | | Initial run | | Rerun 1 | | Rerun 2 | | Rerun 3 | | Rerun 4 | |
|---|---|---|---|---|---|---|---|---|---|---|---|
| Predicted state | | Closed | Open | Closed | Open | Closed | Open | Closed | Open | Closed | Open |
| RMSD to closed crystal[1] (Å) | All-atom | 2.09 | 2.51 | 2.02 | 2.57 | 1.80 | 2.36 | 1.96 | 2.39 | 1.9 | 2.61 |
| | TMD[3] | 0.78 | 1.72 | 0.59 | 1.72 | 0.67 | 1.69 | 0.73 | 1.65 | 0.61 | 1.68 |
| RMSD to open crystal[2] (Å) | All-atom | 2.02 | 1.59 | 2.13 | 1.63 | 2.63 | 1.77 | 2.45 | 1.87 | 2.38 | 1.80 |
| | TMD[3] | 1.56 | 0.68 | 1.67 | 0.66 | 1.68 | 0.67 | 1.63 | 0.77 | 1.64 | 0.76 |

[a]PDB ID 4NPQ
[b]PDB ID 4HFI
[c]Transmembrane domain

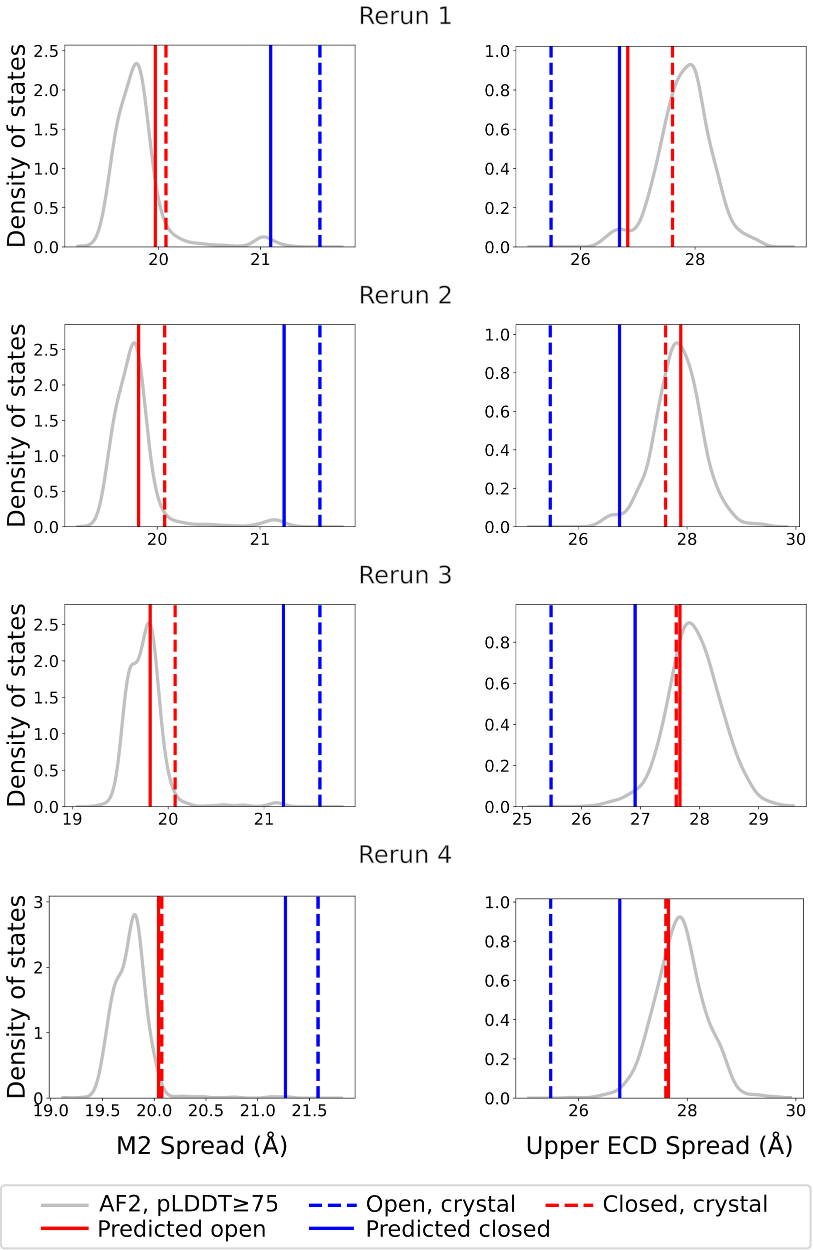


**S8 Fig. M2 spread and upper ECD spread for the reruns.** Distance between the centers of mass of the pore and that of the upper part of the pore lining M2 helix (M2 spread) and the upper spread of the extracellular domain (upper ECD spread) for the predicted structures, the crystal structures, as well as the density of states for all AF2-generated conformations with an average pLDDT score above 75. All data is shown for the four reruns in which distinct functional states were predicted.

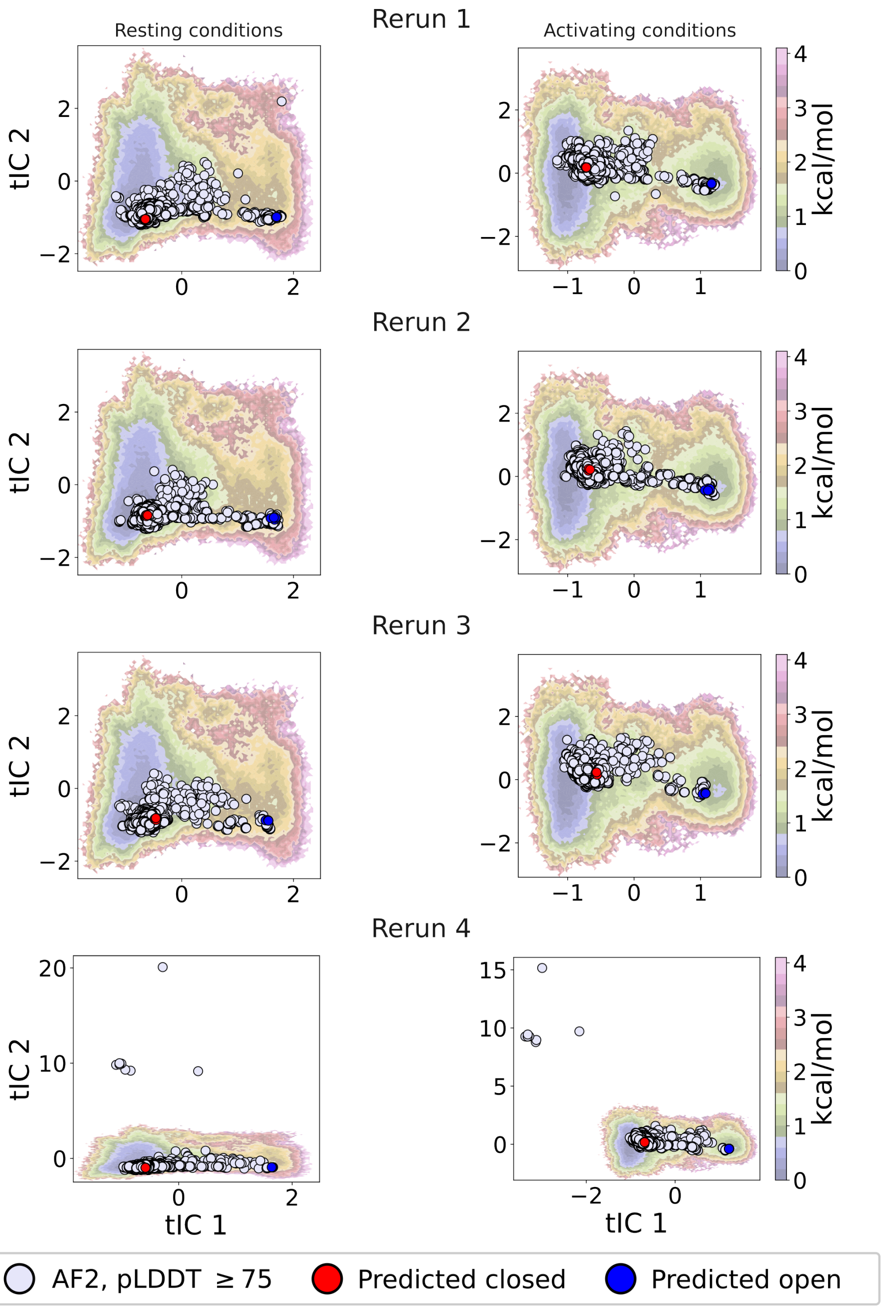


**S9 Fig. Free energy landscape projections for selected conformations.** Projections of the AF2-generated conformations with pLDDT $\geq$ 75 onto the free energy landscape of GLIC at resting and activating conditions, for the four reruns in which distinct functional states were predicted.
